# Long-term evolution of quantitative traits in the *Drosophila melanogaster* species subgroup

**DOI:** 10.1101/2021.11.10.466920

**Authors:** Amir Yassin, Nelly Gidaszewski, Vincent Debat, Jean R. David

**Author notes:** Deceased in 2021 (Jean R. David).

## Abstract

Quantitative genetics aims at untangling the genetic and environmental effects on phenotypic variation. Trait heritability, which summarizes the relative importance of genetic effects, is estimated at the intraspecific level, but theory predicts that heritability could influence long-term evolution of quantitative traits. The phylogenetic signal concept bears resemblance to heritability and it has often been called species-level heritability. Under certain conditions, such as trait neutrality or contribution to phylogenesis, within-species heritability and between-species phylogenetic signal should be correlated. Here, we investigate the potential relationship between these two concepts by examining the evolution of multiple morphological traits for which heritability has been estimated in *Drosophila melanogaster*. Specifically, we analysed 42 morphological traits in both sexes on a phylogeny inferred from 22 nuclear genes for nine species of the *melanogaster* subgroup. We used Pagel’s λ as a measurement of phylogenetic signal because it is the least influenced by the number of analysed taxa. Pigmentation traits showed the strongest concordance with the phylogeny, but no correlation was found between phylogenetic signal and heritability estimates mined from the literature. We obtained data for multiple climatic variables inferred from the geographical distribution of each species. Phylogenetic regression of quantitative traits on climatic variables showed a significantly positive correlation with heritability. Convergent selection, the response to which depends on the trait heritability, may have led to the null association between phylogenetic signal and heritability for morphological traits in *Drosophila*. We discuss the possible causes of discrepancy between both statistics and caution against their confusion in evolutionary biology.

## Introduction

Heritability is a central concept in genetics with crucial applications in medicine and agriculture (Visscher *et al*., 2008). It refers to the tendency of genetically related individuals (e.g., parent-offsprings, sibs, etc.) to resemble each other for a particular phenotype more than they do with other individuals randomly drawn from the population, and that regardless to environmental conditions. The heritability estimate for a trait is a strong indicator for the response of that trait to selection (Falconer, 1960; Lynch & Walsh, 1998). Consequently, long-term parallel selective regimes in different populations could lead to similarity in phenotypic values for a heritable trait among genetically-distant individuals, *i.e*. convergent evolution, due to selection on either independent or shared alleles (e.g., Gross *et al*., 2009; Jones *et al*., 2012; Ralph & Coop, 2015). Therefore, heritability is usually measured within populations using a limited number of generations, where relatedness could be traced and experimental conditions could be controlled.

In spite of such experimental limitations, the concept of heritability has been extended to comparative approaches studying trait evolution across species, where it has mostly been measured in term of phylogenetic signal and was called species-level or phylogenetic heritability (e.g., (Jablonski, 1987; Lynch, 1991; Blomberg *et al*., 2003; Queiroz & Ashton, 2004; Waldron, 2007; Cardini & Elton, 2008; Shirreff *et al*., 2013; de Villemereuil & Nakagawa, 2014; Vrancken *et al*., 2015; Houle *et al*., 2017; Fraser *et al*., 2018; Scher *et al*., 2020). In phylogenetics, a trait with a phylogenetic signal is a trait showing a tendency of closely-related species to resemble each other more than to other species randomly drawn from the phylogeny (Pagel, 1994; Revell *et al*., 2008; Münkemüller *et al*., 2012), a definition that closely resembles that of heritability given above, species playing the role of individuals. However, unlike heritability, phylogenetic signal is by definition measured in situations where convergent selection or genetic introgression could not be controlled, and its biological interpretation has long been problematic (Losos, 2011). Consequently, the relationship between short-term quantitative genetics estimates of heritability and long-term phylogenetic signals remains unclear. Indeed, a positive correlation is expected for neutrally evolving traits, but the relationship is more complex for traits related to fitness.

For nearly a century, *Drosophila melanogaster* has been a quantitative genetics paradigm (Roff & Mousseau, 1987; Mackay, 2010). At the morphological level, four trait categories, namely size, shape, setation (bristle number) and pigmentation have been thoroughly studied on several anatomical structures (e.g., wing, thorax, different abdominal segments, etc.). This has resulted into a wealth of data on heritability, modularity, sexual dimorphism, phenotypic plasticity, response to selection and geographical variation of those traits (reviewed in (Gibert *et al*., 2004). With the advent of genome-wide association techniques, the genomic basis of many of traits is now being unraveled (Mackay & Lyman, 2005; Turner *et al*., 2011; Takahashi *et al*., 2012; Bastide *et al*., 2013, 2016; Dembeck *et al*., 2015; Vonesch *et al*., 2016; Lobell *et al*., 2017; Lafuente *et al*., 2018; Pitchers *et al*., 2019). Of these traits, only wing shape has been analysed using geometrical multivariate approaches within *D. melanogaster* (e.g., (Debat *et al*., 2003; Pitchers *et al*., 2013) as well as among the nine species of the *melanogaster* subgroup (Gidaszewski *et al*., 2009; Klingenberg & Gidaszewski, 2010; Houle *et al*., 2017). However, the relationship between short-term heritability and phylogenetic signals in wing shape traits is still unknown, perhaps due to the nature of this character which could convolute homologization of multivariate statistical summaries (e.g., principal components) within and between species.

In the present study, we investigate the evolution of 42 morphological traits related to size, shape, bristle number, pigmentation and ovariole number, in the nine species of the *Drosophila melanogaster* subgroup. Heritability of most of those traits has been estimated in *D. melanogaster* in terms of intraclass correlation (*t*) using the same isofemale line approach (Delpuech *et al*., 1995; Gibert *et al*., 1998, 2008; Karan *et al*., 1999; Araripe *et al*., 2008). Although *t* involves in theory non-additive components, the ratio between *t* and independent *h^2^* estimates of heritability was found in practice to be 0.95, indicating that *t* is strongly related to *h^2^* (David *et al*., 2005). Moreover, *t* estimates for body size and bristle number traits in the distantly-related drosophilid genus *Zaprionus* approaches those inferred for *D. melanogaster* (David *et al*., 2006; Araripe *et al*., 2008), suggesting that heritability estimates in *D. melanogaster* are likely a good indicator for the trait heritability among its closely-related species. We inferred a molecular phylogeny for the nine species using 22 nuclear genes, and estimated the phylogenetic signal of each trait using Pagel’s λ (Pagel, 1994), because this statistic is the least influenced by the number of analysed species (Münkemüller *et al*., 2012). We also obtained data for 14 climatic variables from the geographical distribution of each species. We found that *t* did not correlate with λ, but it correlated with the phylogenetic regression on the climatic variables. Because heritability determines response to selection, our results indicate that convergent evolution has likely shaped long-term evolution of classical quantitative traits in *Drosophila* leading to the dissociation between their within-species heritability estimates and their between-species phylogenetic signal.

## Materials and Methods

### Drosophila strains

With two exceptions, indicated below, we always used mass cultures founded by a set of at least 30 females or isofemale lines. For the two cosmopolitan species, *D. melanogaster* and *D. simulans*, we used two strains, one from a temperate origin, Prunay (France), collected in 2010 and the other from the island of Sao Tomé at a latitude close to the equator, provided by D. Lachaise in 2002. *D. sechellia* was founded in 1985 by mixing 25 isofemale lines. *D. mauritiana* was founded in the same way in 1988. For *D. teissieri*, we used the type strain, established in 1970 from Mount Selinda in Zimbabwe. For *D. yakuba* we used a still older strain, collected in Kunden (Cameroon) in 1967. For *D. santomea*, we used a mass culture given by D. Lachaise and established in 2005 from Sao Tomé. The last two species corresponded to isofemale lines. For *D. erecta*, we used a line collected in 2005 by D. Lachaise in the park of La Lopé (Gabon), and homozygous for the female light abdomen locus. *D. orena* was the only line available so far, collected in the mountains of Cameroon in 1975.

Groups of 10-20 adult flies were isolated on usual cornmeal sugar food and kept there for 3-4 days. They were then introduced into rearing vials containing a high nutrient, killed yeast medium for producing the experimental flies at a temperature of 21°C. Each group of adults was transferred to a fresh vial for at least 3 successive days. With this procedure, population density in a vial was less than 100. After emergence, young adults were transferred to fresh vials and kept at least for 3 days before being measured. Measurements were made with a binocular microscope equipped with an ocular micrometer, on anesthetized flies (*n* = 10 per sex per strain). For anesthesia we used triethylamine (also called flynap), which has the advantage of producing a long lasting anesthesia on immobile flies.

### Traits measured

A total of 20 different traits were measured in females and 15 in males. They correspond either to linear or area measures (metric traits) or to numbers (meristic traits). Also 4 morphological indices or ratios, were considered in both sexes. These traits were measured either on the thorax or on the abdomen.

#### Body size traits

Three linear measurements were taken, and the dimensions transformed into 100x mm: wing length (WL), from the thoracic articulation to the tip of the wing; thorax length (TL) from the neck to the tip of the scutellum, on a left side lateral view; and thorax width (TW), as the distance between the two posterior large sternopleural bristles, from a ventral view.

#### Body shape indices

From the above data we calculated, on each fly, three ratios: WL/TL which is inversely proportional to the wing loading (Pétavy *et al*., 1997); TL/TW which is the thorax elongation index; and WL/TW which also provides information on wing elongation.

#### Bristles number

Bristles were counted on the thorax and on the ventral side of the abdomen. Thoracic, sternopleural bristles were counted on each side and the sum (STPn) was used in calculations. We also measured the length of the two major bristles, anterior (STPa) and posterior (STPp) sternopleurals, and also calculated their ratio (STPp/a). The inverse of this ratio, called spernopleural index (Sturtevant, 1942), is often used in systematics. Abdominal bristles were counted on the successive sternites, from segment 2 up to segment 5 in males (A2-A5) and segment 7 in females (A2-A7). The bristles were also counted on each side of tergite 7 in females, and the sum calculated (T7).

#### Abdominal pigmentation

This was measured on successive tergites (P2 to P7 in females and P2 to P6 in males). The surface of black area at the posterior part of each tergite was visually scored by establishing 11 phenotypic classes from 0 (no black pigment) up to 10 (tergite completely black) (David *et al*., 1990).

#### Ovariole number

Mature females were dissected and the ovariole number counted on each ovary and then the sum calculated (OV).

### Phylogeny inference

We chose the 22 nuclear genes that were analysed by (Akashi *et al*., 2006) to investigate codon usage evolution in five species of the *melanogaster* subgroup: *D. melanogaster*, *D. simulans*, *D. teissieri*, *D. erecta* and *D. orena*. We verified sequences of these genes in the five species of the subgroup with published, annotated complete genome sequence: *D. melanogaster*, *D. simulans*, *D. sechellia*, *D. yakuba* and *D. erecta* (Clark *et al*., 2007). We also obtained their DNA sequences from the published genomes of *D. mauritiana* (Nolte *et al*., 2013) and *D. santomea* (Turissini *et al*., 2015) using BLAST (Altschul *et al*., 1997). Sequences from each gene were separately aligned with Muscle (Edgar, 2004) as implemented in the MEGA 7.0 software package (Kumar *et al*., 2016). We also used MEGA to estimate the best substitution model for each gene, which were HKY + Γ for *Adh*, *Adhr*, *Amyrel*, *AP-2mu*, *boss*, *dpp*, *Fur2*, *g*, *Gld*, *Gpdh*, *Mcm3*, *pav*, *per*, *RpII215*, *run*, *twi* and *Zw*, HKY + I for *Amy-p* and *Cyp28c1*, GTR + Γ for *LanB2* and *ry*, and HKY + Γ + I for *Osbp*. We concatenated the genes (41,382 nucleotides) and inferred a Bayesian phylogeny using MrBayes v3.2.4 (Ronquist *et al*., 2012), applying the estimated substitution model to each gene partition. Two runs of 500,000 generations were conducted and sampled every 1,000 generations under a strict clock model. We assessed convergence – the average standard deviation of split frequency < 0.01 and the potential scale reduction factor ~1.00 – using MrBayes. A burn-in period of 25% of samples was used. Alignment and MrBayes commands are given in Supplementary Text 1.

### Phylogenetic signal

Morphometric data were analysed as univariate variables to infer the mean and standard error per species in R (www.r-project.org). R was also used to conduct Principal Component Analysis (PCA) on the complete (multivariate) data set after excluding ratio traits. Mean values were mapped on the species phylogeny (excluding data of temperate strains of *D. melanogaster* and *D. simulans*). We inferred phylogenetic signal (species-level heritability) in terms of (Pagel, 1994) lambda (λ). We used the Generalized Least Square (GLS) approach to infer λ as implemented in the Continuous Model A method assuming random walk in the BayesTraits v2.0 software package (Pagel & Meade, 2014). All analyses were conducted using MCMC chains of 1,010,000 iterations and 1,000 post-burnin trees on trees generated by MrBayes to account for topological uncertainties. The harmonic mean of the likelihoods, *i.e*., marginal likelihoods of the MCMC runs, was checked for stability. We estimated λ significance in terms of Bayes factors (BFs) by comparisons with runs where λ was restricted to 0 (*i.e*. the trait evolution is independent from the phylogeny) or 1 (*i.e*. the phylogeny correctly predicts the trait evolution). BF was estimated as 2 * (marginal likelihood of the best-fit model – marginal likelihood of the worse-fit model). We considered BF > 2, 5 or 10 as supportive, strong, or very strong support of the best-fit model, respectively, following the general convention (Pagel & Meade, 2014).

In order to examine the relationship between phylogenetic signal (λ) and short-term heritability, we mined the quantitative genetics literature in *D. melanogaster* to obtain heritability estimates for the morphometrical traits. Because heritability could be estimated using various approaches, we opted for the sake of comparison for a single methodology, *i.e*. intra-class correlation (*t*), that has been used in most of the traits analysed here (Delpuech *et al*., 1995; Gibert *et al*., 1998, 2008; Karan *et al*., 1999; Araripe *et al*., 2008). To the best of our knowledge, no heritability estimates using *t* was made for any of the analysed traits in other species of the *melanogaster* subgroup. However, *t* estimates for body size and bristle number traits in the distantly-related drosophilid genus *Zaprionus* approaches those inferred for *D. melanogaster* (David *et al*., 2006; Araripe *et al*., 2008), suggesting that heritability estimates in *D. melanogaster* are likely a good indicator for the trait heritability among its closely-related species. Therefore, we inferred the coefficient of correlation between λ and *t*, as an indication for the relationship between short-term heritability and long-term phylogenetic signal.

### Phylogenetic regression on climatic variables

In order to determine which environmental variables might also explain the evolution of each trait, we conducted a regression analysis for each trait (except for shape ratios) on species means of 14 climatic variables. These included seven climatic sets (namely, average (Tavg), minimum (Tmin) and maximum (Tmax) air temperatures and earth temperature (TEarth) measured in °C, relative humidity (RH) in %, atmospheric pressure (AP) in kPa, and wind speed (WS) at 50 m above ground in m/s) and seven radiation sets (namely, insolation incident (II), diffuse radiation (DR), direct normal radiation (DNR) and latitude tilt radiation (LTR), each measured in kWh/m^2^/day, clear sky (CS) in number of days with no clouds, no sun (NS) in number of days with clouds, and ultra-violet index (UVI) in units of 25 mW/m^2^). We first obtained geographical coordinates (*i.e*. latitudes and longitudes) from TaxoDros v1.04 (Bächli 2016; last accessed December 2020) for each locality from which each of the nine species was collected in tropical Africa (*i.e*. south to latitude 17°N). We then extracted for each locality climatic data averaged over 22 years (from 1983 to 2005) as monthly averages from the NASA Surface meteorology and Solar Energy: Global Data Sets website (www.eosweb.larc.nasa.gov) for all variables, except UV index data which were from the Tropospheric Emission Monitoring Internet Service (www.temis.nl) as average values of the year 1999. For all variables, we estimated the annual average per geographical locality, and for each species over all localities. We mapped mean values on the species phylogeny and estimated their phylogenetic signal as for morphometrical traits using BayesTraits. We then estimated the coefficient of phylogenetic regression (β) of each morphological trait on each climatic variable using the Continuous model in BayesTraits to determine the sign of the relationship, and estimated its significance by counting the ratio of the time β crossed the zero point. A well-supported β should minimally switch signs.

In order to be able to obtain an estimate of the “overall effect” of the climatic factors on each trait, we conducted a PCA analysis on the average values of the 14 climatic variables for each species, and then tested the phylogenetic regression using BayesTraits for each quantitative trait and values of the two first principal components, henceforth estimated two scores, denoted ß_PC1_ and ß_PC2_, respectively. We then tested the correlation between the absolute values of the regression coefficients, *i.e*. |ß_PC1_| and |ß_PC2_|, with both *t* and λ for each morphological trait.

## Results

### *Multivariate analysis of morphological traits recovers the two major clades of the* melanogaster *subgroup*

The Bayesian phylogeny of the nine species inferred from the 22 nuclear genes (Figure 1A) recapitulates the current view of relationships based on complete genome sequences of the five annotated species (Clark *et al*., 2007), with all nodes having a posterior probability of 100%. The subgroup is divided into two major clades: *melanogaster* (4 species) and *yakuba* (5 species). Principal component analysis (PCA) of the 20 traits in the 11 strains showed a strong concordance with the *melanogaster-yakuba* phylogenetic split for both females and males (Figure 1B,C). The first four principal components explained 69.5% and 76.1% of the total variance in females and males, respectively. PC1 (34.3% and 34.8% in females and males, respectively) was mostly correlated with abdominal pigmentation traits, PC2 (17.2% and 18.3%) with abdominal bristles in both sexes and ovariole number in females, PC3 (9.8% and 14.5%) with body size, and PC4 (8.2% and 8.5%) with thoracic bristles.

**Figure 1.**
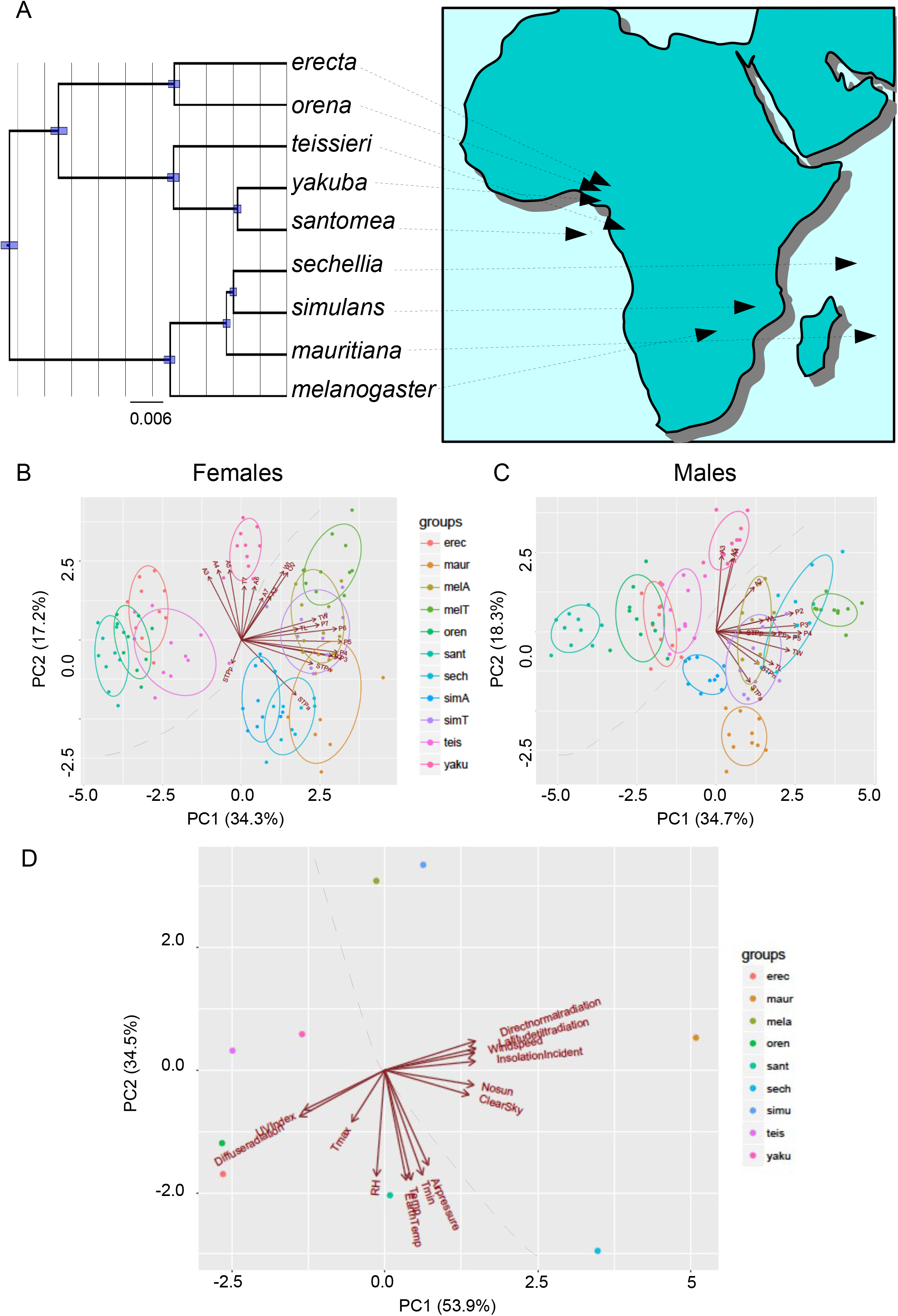
(A) Bayesian phylogeny of the nine species of the *melanogaster* subgroup inferred from 22 nuclear genes with arrows indicating on the map median geographical coordinates for each species (according to data fromTAXODROS, see text). (B,C) Principal component analysis (PCA) of female (B) and male (C) morphological data. (D) PCA of averaged values per species of 14 climatic variables. Dashed line indicates the difference between the *melanogaster* and *yakuba* clades.

On average (Table S1), the abdomen of the species of the *melanogaster* clade is three times darker than that of the *yakuba* clade (summed over the six segments). The two temperate populations of the two cosmopolitan species (*D. melanogaster* and *D. simulans*) correctly fall within the morphospace of this clade, indicating the persistence of the pigmentation difference (Figure 1B,C). Within the *yakuba* clade, *D. yakuba* is exceptional in having a very bristly abdomen (summed over the six segments) and a dark abdomen comparable to those of the *melanogaster* clade (Figure 1B,C). The completely yellow abdomen of *D. santomea* males (Lachaise *et al*., 2000) clearly distinguishes it from the remaining species of the subgroup where the last two segments of the male abdomen are completely pigmented (Figure 1C).

### Phylogenetic signal of morphological traits does not correlate with short-term heritability

Estimates of the phylogenetic signal (λ) revealed heterogeneity of the effect of the phylogenetic history among trait categories (Table 1). On average over traits and sexes, λ ranged from 0.37 to 0.43 for body size, shape and ovariole number, and from 0.51 to 0.52 for setation and pigmentation traits. There was no correlation between λ and heritability as estimated from the intraclass correlation in *D. melanogaster* (*r* = −0.035, *P* = 0.860) after combining data from Table 1 from both sexes.

**Table 1.** Phylogenetic signal (λ), heritability as intra-class correlation among isofemale lines (*t*) and average absolute coefficient of phylogenetic regression |β| of PC1 and PC2 scores of climatic variables for 44 morphological traits in the nine species of the *Drosophila melanogaster* species subgroup. Bold values correspond to averages per trait categories.

| Trait | Females |  |  |  | Males |  |  |  |
| --- | --- | --- | --- | --- | --- | --- | --- | --- |
| | $t$ | $\lambda$ | $\vert \beta_{PC1}\vert $ | $\vert \beta_{PC2}\vert $ | $t$ | $\lambda$ | $\vert \beta_{PC1}\vert $ | $\vert \beta_{PC2}\vert $ |
| **Body size** |  |  |  |  |  |  |  |  |
| WL | 0.50 | 0.40 | 0.82 | 0.98 | 0.44 | 0.44 | 0.77 | 0.96 |
| TL | 0.38 | 0.33 | 0.92 | 0.56 | 0.29 | 0.38 | 0.96 | 0.68 |
| TW |  | 0.43 | 0.92 | 0.91 |  | 0.46 | 0.94 | 0.96 |
| **Body shape** |  |  |  |  |  |  |  |  |
| WL/TL | 0.37 | 0.46 |  |  | 0.37 | 0.44 |  |  |
| WL/TW |  | 0.48 |  |  |  | 0.47 |  |  |
| TL/TW |  | 0.36 |  |  |  | 0.37 |  |  |
| **Setation** |  |  |  |  |  |  |  |  |
| STPn | 0.31 | 0.73 | 0.92 | 0.52 | 0.28 | 0.53 | 0.77 | 0.92 |
| STPa |  | 0.76 | 0.99 | 0.75 |  | 0.61 | 0.96 | 0.92 |
| STPp |  | 0.37 | 0.91 | 0.98 |  | 0.38 | 0.94 | 0.96 |
| STPr |  | 0.79 |  |  |  | 0.62 |  |  |
| A2 | 0.11 | 0.31 | 0.74 | 0.73 | 0.21 | 0.32 | 0.82 | 0.71 |
| A3 | 0.19 | 0.60 | 0.79 | 0.52 | 0.23 | 0.42 | 0.64 | 0.89 |
| A4 | 0.23 | 0.63 | 0.68 | 0.61 | 0.15 | 0.45 | 0.72 | 0.68 |
| A5 | 0.12 | 0.64 | 0.63 | 0.51 | 0.06 | 0.40 | 0.75 | 0.63 |
| A6 | 0.09 | 0.36 | 0.59 | 0.57 |  |  |  |  |
| A7 | 0.17 | 0.37 | 0.76 | 0.68 |  |  |  |  |
| T7 |  | 0.46 | 0.97 | 0.89 |  |  |  |  |
| **Pigmentation** |  |  |  |  |  |  |  |  |
| P2 | 0.26 | 0.65 | 0.75 | 0.57 | 0.26 | 0.42 | 0.58 | 0.74 |
| P3 | 0.21 | 0.69 | 0.70 | 0.57 | 0.28 | 0.51 | 0.50 | 0.64 |
| P4 | 0.22 | 0.68 | 0.60 | 0.53 | 0.23 | 0.54 | 0.65 | 0.69 |
| P5 | 0.36 | 0.62 | 0.71 | 0.71 |  | 0.40 | 0.75 | 0.81 |
| P6 | 0.41 | 0.49 | 0.55 | 0.92 |  | 0.44 | 0.77 | 0.83 |
| P7 | 0.33 | 0.41 | 0.65 | 0.99 |  |  |  |  |
| **Reproductive organs** |  |  |  |  |  |  |  |  |
| OV | 0.37 | 0.37 | 0.88 | 0.99 |  |  |  |  |

### Phylogenetic regression reveals a substantial effect of climatic variables on morphological evolution

Principal component analysis (PCA) of the average values of 14 climatic variables deduced from biogeographical data of the nine species of the *melanogaster* subgroup revealed a strong distinction between the two major clades (Figure 1D). The distinction was most obvious along PC1, which explained 54% of the variance and mostly correlated with solar radiation variables in agreement with the East-West disjunction of the likely origin of the two clades. PC2, however, which explained 35% of the variance, was most correlated with temperature and relative humidity variables. It mostly discriminated among the most closely-related species, *i.e. D. simulans* from both *D. mauritiana* and *D. sechellia*, and *D. yakuba* from *D. santomea*.

Phylogenetic signals (λ) for species means of climatic variables fell within two categories: low (λ = 0.32-0.43) for temperature, air pressure and relative humidity variables, and intermediate to high (λ = 0.53-0.74) for wind speed and radiation variables (Table S2). We found some correspondences between λ estimates for morphological traits and climatic variables and their phylogenetic regressions (ß) (Figure 2). For example, morphological traits with low λ, such as wing length, ovariole number or pigmentation of the last abdominal segment in females (P7) were significantly negatively correlated with climatic variables with low λ such as temperature and relative humidity.

**Figure 2.**
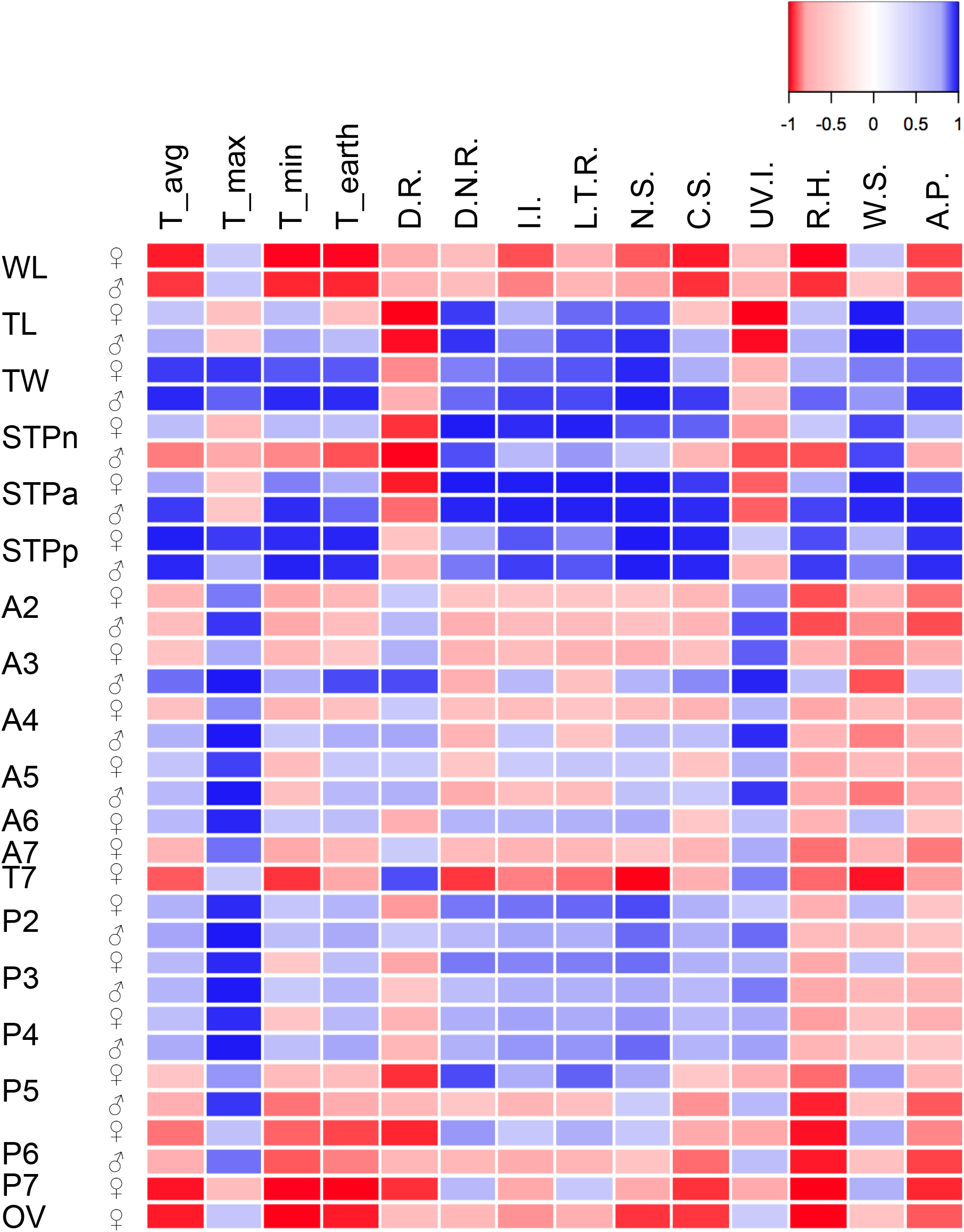
Heatmap of (1 – *P*) where *P* is the significance probability of the phylogenetic regression of each phenotypic trait on 14 ecological variables, signed according to the coefficient of regression.

Because ß varied substantially among individual climatic variables, and climatic variables are usually correlated, we conducted the phylogenetic regression analyses on the PC1 and PC2 scores of the species climatic data (Table 1). For all traits and sexes (combined from Table 1), |ß_PC1_| did not correlate with either *t* (*r* = −0.248, *P* = 0.221) or λ (*r* = −0.070, *P* = 0.691). On the contrary, |ß_PC2_| had a highly significant positive correlation with *t* (*r* = 0.585, *P* = 1.686 x 10^−3^) and negatively correlated with λ (*r* = −0.421, *P* = 0.012).

In summary, short-term heritability estimates do not correlate with phylogenetic signal, *i.e*. the so-called species-level or phylogenetic heritability, but they do correlate with phylogenetic regression on species-averaged climatic variables related to temperature and aridity.

## Discussion

In spite of their conceptual resemblance, *i.e*. increased similarity due to genetic relatedness, our results do not support a correlation between the measurements of heritability and phylogenetic signal, at least for quantitative morphological traits in *Drosophila*. On the one hand, heritability is explained by the genetic architecture of a trait, whereas phylogenetic signal refers to how evolution in one trait correlates with speciational history. Low phylogenetic signal indicates high homoplasy, which could be caused by a multitude of evolutionary factors, dependent on or independent from the genetic architecture of the trait. Our observation that quantitative morphological characters in *D. melanogaster* show a significant phylogenetic regression on climatic variables (Figure 2), suggests that convergent selection on those heritable traits might have been the cause of their low phylogenetic signal, *i.e*. homoplasy.

Body size, bristle number and pigmentation have been classical traits in *Drosophila* quantitative genetics for nearly a century, but they have rarely been used in systematics or phylogenetics due to their low phylogenetic signal (Al Sayad & Yassin, 2019). In fact, even within *D. melanogaster*, most of those characters show evidence for convergent evolution among geographical populations inhabiting colder regions (e.g., high altitudes and latitudes on different continents) (Gibert *et al*., 2004). For wing size for example, larger wings in high latitude populations in Australia and South America are mostly driven by different cellular mechanisms, namely cell number proliferation and cell size enlargement, respectively (Zwaan *et al*., 2000). In high altitude Ethiopian populations with the largest size traits in *D. melanogaster*, cell size enlargement was also found to significantly drive the evolution of larger wings (Lack *et al*., 2016b; c), in spite of the fact that those flies are genetically closer to normal size African populations than to other non-African flies (Lack *et al*., 2016a). For pigmentation, convergent evolution of melanism occurs between high latitude populations and high altitude African populations, but the underlying genetic basis differs between populations (Bastide *et al*., 2014, 2016). Even when the same genetic locus underlies the same trait within and between species, apparent phylogenetic similarities can still hide distinct genetic origins, such as the light female morph phenotype caused by independent mutations at the same regulatory element in the sister species *D. erecta* and *D. orena* (Yassin *et al*., 2016).

Our results might have been biased by our choice of characters. Indeed, an analysis of the phylogenetic signal of 490 morphological characters in drosophilids revealed that nearly two thirds of the characters evolve convergently (Al Sayad & Yassin, 2019). However, genital traits were among those with the highest signal. In the *melanogaster* subgroup, male and female genital structures are indeed the most discriminatory characters between species (Yassin & Orgogozo, 2013). Due to their small size and dissection-associated difficulties, few studies have aimed at investigating their quantitative genetics. It was however estimated that the heritability of the size of the male epandrial posterior lobe, which is used to grasp the female during copulation, was 0.65 in *D. melanogaster* (Takahara & Takahashi, 2015) and as high as 0.93 in crosses between *D. simulans* and *D. mauritiana* (Zeng *et al*., 2000). Here, a correspondence between high heritability and phylogenetic signal for the same trait may be expected, if the trait evolution has contributed to the phylogenetic divergence between species.

In conclusion, our results caution against the usage of the term heritability for phylogenetic signal, even if some precisions such as phylogenetic or species-level heritability are applied. Phylogenetic signal is a description of a pattern that can be generated by a multitude of processes (Losos, 2011). Although a correspondence between the two concepts may sometimes be found, mostly at short time scales (e.g., in viruses, (Shirreff *et al*., 2013; Vrancken *et al*., 2015), it is most likely, given the preponderance of homoplasy for most types of characters, that a strict correlation will be rare. Our results also demonstrate a potential role of environmental conditions in shaping the genetic architecture of a trait. The positive correlation between short-term heritability and long-term phylogenetic regression on variables related to temperature and aridity may be due to the fact intraclass correlations estimates of heritability may either involve a part of the genotype x environmental variance (*i.e*. broad-sense heritability) or to the direct action of selection on the genetic variance underlying quantitative traits. Future dissection of the genetic architecture of the traits analysed here in the remaining species of the *melanogaster* subgroup would definitively help explaining the role of the environment in the long-term evolution of those classical *Drosophila* quantitative traits.

## Conflicts of interest

We declare no conflict of interest.

## Notes

### Competing Interest Statement

The authors have declared no competing interest.

